# Collateral connectomes of *Esr1*-positive hypothalamic neurons modulate defensive behavior plasticity

**DOI:** 10.1101/2025.01.10.632334

**Authors:** Veronika Csillag, Chiara Forastieri, Gréta Martina Szücs, Inés Talaya Vidal, Marianne Hiriart Bizzozzero, Luke D. Lavis, Daniela Calvigioni, János Fuzik

**Affiliations:** Department of Neuroscience, Karolinska Institutet, Stockholm, Sweden; Janelia Research Campus, Howard Hughes Medical Institute, Ashburn, VA, USA

## Abstract

The ventromedial hypothalamus (VMH) projects to the periaqueductal gray (PAG) and anterior hypothalamic nucleus (AHN), mediating freezing and escape behaviors, respectively. We investigated VMH collateral (VMH-coll) neurons, which innervate both PAG and AHN, to elucidate their role in postsynaptic processing and defensive behavior plasticity. Using all-optical voltage imaging of 22,151 postsynaptic neurons *ex vivo*, we found that VMH-coll neurons engage inhibitory mechanisms at both synaptic ends and can induce synaptic circuit plasticity. *In vivo* optogenetic activation of the VMH-coll somas induced escape behaviors. We identified an *Esr1*-expressing VMH-coll subpopulation with postsynaptic connectome resembling that of wild-type collaterals on the PAG side. Activation of Esr1+VMH-coll neurons evoked freezing and unexpected flattening behavior, previously not linked to the VMH. Neuropeptides such as PACAP and dynorphin modulated both Esr1+VMH-coll connectomes. In vivo κ-opioid receptor antagonism impaired Esr1+VMH-coll-mediated defensive behaviors. These findings unveiled the central role of VMH-coll pathways in innate defensive behavior plasticity.

## Main

Defensive fight-or-flight behaviors are fundamental survival strategies enabling animals to evade predation or escape life-threatening situations. The ventromedial hypothalamus (VMH), a key component of the innate defense circuitry, topographically encodes olfactory threat stimuli^1,2^. The VMH connects to multiple regions, that mediate distinct defensive responses, including the periaqueductal gray matter (PAG) that induces freezing^3,4^, the anterior hypothalamic nucleus (AHN), which integrates predator-proximity cues to regulate defensive behaviors^5,6^ and the premammillary nucleus (PMd) which processes predatory threats^7,8^. Steroidogenic factor-1 (SF1) expression delineates the defensive subdivision of the VMH^9^. Optogenetic activation of SF-1 positive VMH neurons have been shown to induce biphasic defensive behavior initiating with freezing and transitioning to intense escape responses^10^. Independent activation of SF1-expressing VMH-PAG and VMH-AHN pathways mediates opposing defensive behaviors, driving freezing and promoting escape, respectively^11^. Although collateral neurons within the VMH that partially overlap with SF1 expression have been identified, their identity, synaptic connectivity, and functional role in modulating the flexibility of defensive behaviors remain unknown.

The flexible adjustment of innate behaviors is fundamental to survival^12,13^. Central to this adaptability is plasticity that reorganize synaptic circuits to fine-tune behavioral responses. Neuropeptides released during heightened neuronal activity^14^ play a pivotal role in shaping the functional connectivity of synaptic networks. The interplay between fast synaptic transmission and neuropeptide co-transmission orchestrates large-scale circuit adaptations that drive behavioral responses and influence chronic brain states^15,16^. Within the ventromedial hypothalamus (VMH), neuropeptides such as Substance P (SP), PACAP, and dynorphin modulate defensive and anxiety-related responses^17–21^. Notably, dynorphin has been implicated in stress-induced dysphoria via the κ-opioid receptor (KOR) system^22,23^. However, our understanding of how neuromodulation shapes postsynaptic processing within the innate defense circuitry and its impact on behavioral plasticity remains limited.

In the present study, we investigated the role of collateral VMH (VMH-coll) neurons in modulating PAG and AHN postsynaptic processing and defensive behaviors. We found that VMH-coll neurons establish more pronounced synaptic connectivity with the AHN compared to the PAG. Interestingly, both postsynaptic targets engaged GABAergic inhibitory synaptic transmission that persistently diminished following optical high-frequency stimulation (oHFS) of both collateral ends. Despite detecting fewer synaptic responses at the VMH-PAG-coll terminal, we demonstrated that its activation induced robust freezing behavior, whereas somatic activation of VMH-coll neurons triggered escape behaviors such as jumping, without a phase of immobility. Repeated activation of VMH-coll somas resulted in behavioral plasticity with more pronounced escape and with reversed innate defensive responses to predator scent, shifting immobility into escape behaviors. We identified a subpopulation of VMH-coll neurons expressing the *Esr1* gene and revealed that their postsynaptic response profile closely resembled that of the VMH-PAG-coll connectome, with more distinct differences from the VMH-AHN-coll circuitry. In line with this, we found that somatic activation of Esr1-VMH-coll neurons elicited freezing and flattening behaviors characteristic of a hiding response. We also showed that both synaptic ends of the Esr1-VMH-coll connectome were modulated by neuropeptides, including PACAP and dynorphin, which exerted overall excitatory effects. Remarkably, we found that *in vivo* optogenetic activation of Esr1-VMH-coll neurons under systemic KOR antagonism disrupted behavioral flexibility by reducing the potentiation of immobility, impairing the adaptability of the defensive behavior. Collectively, our findings revealed that dynorphin-mediated neuromodulation is indispensable for the behavioral flexibility of innate defensive responses driven by the Esr1-VMH-coll neuronal population.

## Results

### Anatomical organization of the VMH-coll pathway

To map the anatomy of the VMH-coll pathway we visualized the VMH-PAG and the VMH-AHN neuronal populations using retrograde viral labeling (Fig. 1a). We found that VMH-coll neurons constitute approximately 25% of the total AHN and PAG innervating VMH, where VMH-AHN projections gave ∼30%, and VMH-PAG projections gave ∼45% of VMH-PAG projections (Fig. 1b,c). To resolve the spatial distribution of VMH-coll neurons within the VMH, we performed a three-dimensional mapping of VMH-coll neurons in four mice using the Allen Common Coordinate Framework^24^(Fig. 1d). We found that ∼40-50% of the VMH-coll neurons were in the dorso-medial VMH (dmVMH), ∼35% were in the central VMH (cVMH), primarily in the posterior VMH, and about 10-15% of VMH-coll neurons resided in the ventro-lateral VMH (vlVMH) (Fig, 1e). We also found that the VMH-coll neurons form robust axon terminal coverage in both synaptic ends of the VMH-coll pathway in the PAG and AHN (Fig. 1f), regions involved in driving opposing fight-or-flight defensive behaviors^11^.

**Fig. 1:**
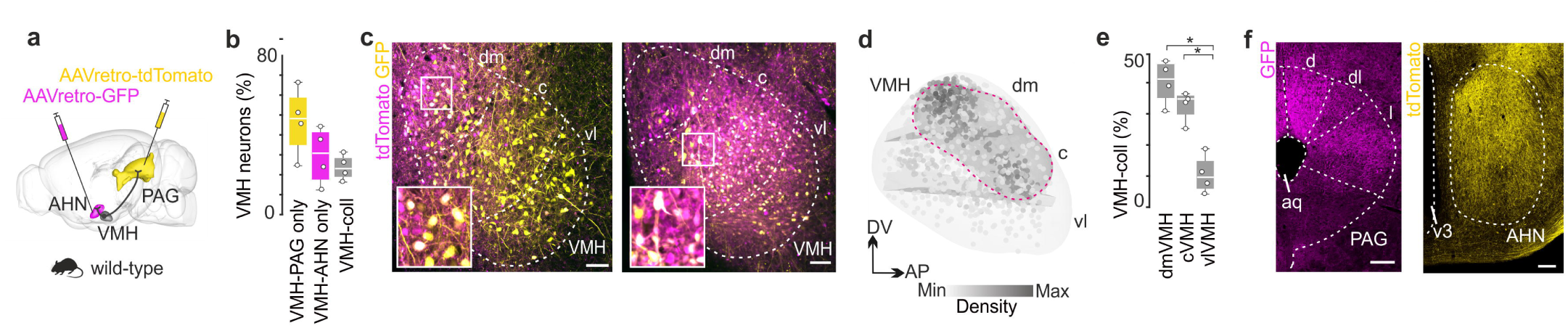
Anatomical organization of the VMH-coll pathway. **a,** Scheme of retrograde viral injections to label and visualize VMH-coll neurons. **b,** Box plot comparing the proportion of only PAG-projecting (yellow), AHN-projecting (purple) and collateral neurons projecting to both PAG and AHN (gray). **c,** Confocal images of PAG-projecting (yellow), AHN-projecting (magenta) and colocalization (white) marking VMH-coll neurons. **d,** Side view of three-dimensional VMH plot with spatial mapping of VMH-coll neurons and the encapsulating volume covering the density core of VMH-coll neurons (gray blob). **e,** Box plot comparing the spatial distribution of VMH-coll neurons in the dmVMH, cVMH and vlVMH. **f,** VMH-coll axonal coverage in the PAG (left) and in the AHN (right).

### Postsynaptic response types of the VMH collateral connectomes

To investigate potential biases in the recruitment of postsynaptic connectomes and differences in postsynaptic response types (PRTs) at the two synaptic ends of VMH-coll neurons, we employed all-optical postsynaptic voltage imaging with the Voltron sensor^25,26^. To achieve systemic Cre-independent expression of somatic-targeted Voltron (Voltron-ST) for large anatomical coverage, we developed a PHP.eB virus delivered via a single retro-orbital injection (Fig. 2a). For *ex vivo* imaging, Voltron-ST was fluorescently labeled with Janelia Fluor®585-HaloTag (JF-585)^27^ (Fig. 2b). Pathway-specific expression of Channelrhodopsin-2 (ChR2) in VMH-PAG-collateral axons was achieved via a dual viral strategy. Retrograde-AAV delivered Cre recombinase to AHN inputs, followed by an anterograde-AAV injection into the VMH to restrict ChR2 expression to VMH-PAG-collateral neurons (Fig. 2a,b). The reverse strategy labeled VMH-AHN-collateral axons (Fig. 2a,b). Ex vivo, we all-optically imaged 7143 neurons across anterior-posterior axes in six fields of view (FOVs; 200 × 500 µm) per serial coronal slice from five mice per region, capturing 3212 neurons in the PAG and 3931 in the AHN (Fig. 2c). PRTs were classified based on extracted optical-physiology (o-phys) traces. Optical action potentials (o-APs), subthreshold (o-Sub) kinetics, excitatory and inhibitory PSPs (o-EPSPs, o-IPSPs), and burst activity were detected in each o-phys trace. Using 29 o-phys parameters, we performed unbiased hierarchical clustering to categorize VMH-coll PRTs. In the VMH-PAG-coll connectome, 3212 PRTs were grouped into 10 clusters (Fig. 2d,e). Excitatory responses ranged from strong excitation inducing burst firing (cluster 1) to mild excitation in spontaneously active (cluster 2) or silent neurons (cluster 5). Inhibitory responses included mild inhibition (clusters 6, 7), pronounced inhibition with rebound activation (cluster 8), and strong inhibition silencing active neurons (cluster 9). Clusters 3 and 10 represented suppressive and weak responses, respectively. Notably, spontaneous AP firing was associated with clusters showing pronounced responses. The VMH-AHN-coll connectome exhibited greater diversity, with 3931 PRTs categorized into 13 clusters (Fig. 2f,g). Excitatory responses included persistent activity (cluster 3), time-locked APs (cluster 2), and burst firing (cluster 5). Inhibitory responses ranged from rebound bursting (clusters 9, 11) to strong inhibition silencing active neurons (cluster 12). Notably, AHN connectivity was detected in 75% of probed neurons, with 38% showing suprathreshold or strong inhibitory responses regulating AP firing. By contrast, PAG connectivity was found in only 50% of neurons, with 7% receiving strong synaptic input capable of driving AP firing. This is a fraction of the synaptic drive of the complete VMH-PAG connectome, that showed 89% connectivity where all-optical testing evoked AP firing in ∼67% of PAG neurons^26^. Surprisingly, both connectomes exhibited both excitatory and inhibitory PRTs, diverging from the predominantly excitatory glutamatergic profile of the VMH^26^. Collectively, these findings reveal a quantitative bias of synaptic input favouring the VMH-AHN-col pathway and highlight the functional diversity within the VMH-collateral connectomes.

**Fig. 2:**
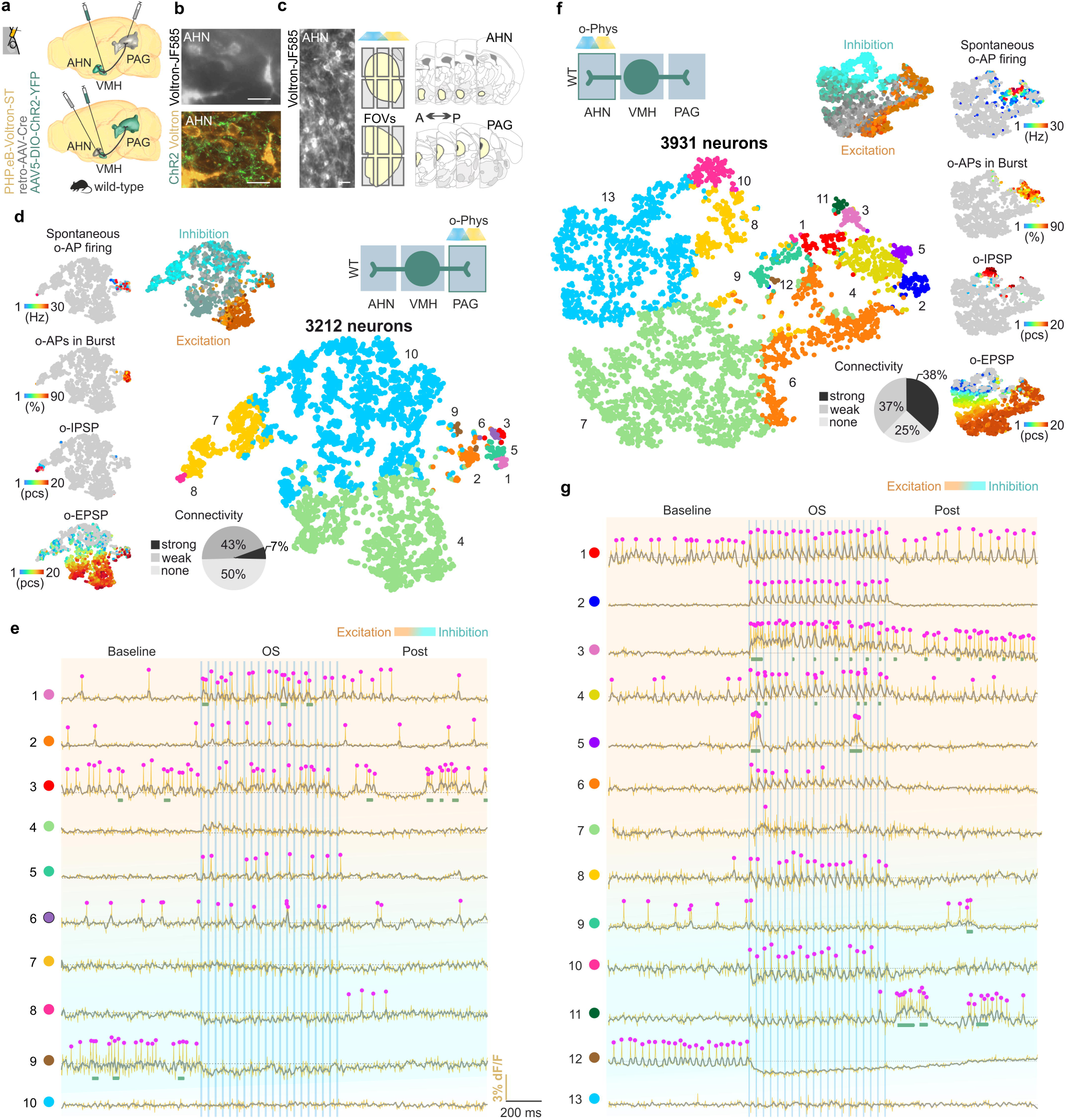
Postsynaptic response types of the VMH collateral connectomes. All Voltron traces were reversed. **a,** Scheme of viral expression of ChR2 in the VMH-AHN-col (top) and VMH-PAG-col (bottom) collateral pathways and systemic expression of Voltron-ST. **b,** Epifluorescent image of neurons with JF-585 signal (top, scale bar, 20Lμm); Confocal image of the same neurons with ChR2 (green) and JF-585-Voltron-ST (gold) labeling in the AHN (bottom; scale bar, 20Lμm). **c,** Example of an AHN FOV with JF-585-Voltron-ST neurons (left, scale bar, 20Lμm) and the illustration of tile-covering AHN (top) and PAG (bottom) for all-optical imaging covering AP coordinates of AHN and PAG (right). **d,** VMH-PAG-col o-phys connectome; t-SNE plot of inhibitory (blue) and excitatory (red) PRTs (top), t-SNE plots of PRTs color coded by Spontaneous o-AP firing, o-APs in Burst, o-IPSP and o-EPSP (left), t-SNE plot of the identified o-phys clusters (right), pie chart showing proportion of no connection (light gray), weak connections (dark gray) and strong connections (black) in the VMH-PAG connectome (bottom). **e,** Segments of all-optical sweeps with Baseline, OS and Post (top). Representative excitatory and inhibitory (top to bottom, right, blue-red bar) o-phys traces of the identified VMH-PAG-col o-phys clusters (color code and number is identical to d), green bars mark bursts, magenta dots mark o-APs, gray dashed line marks baseline voltage, blue bars mark the timing of 20 Hz 473 nm OS. **f,** VMH-AHN-col o-phys connectome; t-SNE plot of inhibitory (blue) and excitatory (red) PRTs (top), t-SNE plots of PRTs color coded by Spontaneous o-AP firing, o-APs in Burst, o-IPSP and o-EPSP (right), t-SNE plot of the identified o-phys clusters (left), pie chart showing proportion of no connection (light gray), weak connections (dark gray) and strong connections (black) in the VMH-PAG connectome (bottom). **g,** Segments of all-optical sweeps with Baseline, OS and Post (top). Representative excitatory and inhibitory (top to bottom, right, blue-red bar) o-phys traces of the identified VMH-PAG-col o-phys clusters (color code and number is identical to d), green bars mark bursts, magenta dots mark o-APs, gray dashed line marks baseline voltage, blue bars mark the timing of 20 Hz 473 nm OS.

### Circuit plasticity of the VMH collateral connectomes

Both AHN and PAG VMH-coll postsynaptic connectomes exhibited excitatory and inhibitory synaptic processing. To investigate how excitatory-inhibitory interplay reshapes the connectome following high levels of pathway activation, we applied optical high-frequency stimulation (oHFS) to synaptic terminals in both the PAG and AHN *ex vivo* and analyzed changes in PRTs using VoltView analysis package^26^ (Fig. 3a) All-optical voltage imaging of the VMH-PAG-coll connectome revealed that responses in cluster 4, characterized by weak excitatory connections, became more depolarized after oHFS. A similar depolarizing shift was observed in inhibitory PRTs of cluster 7. Additionally, we detected a reduction in spontaneous firing activity and a decrease in firing frequency within bursts (Fig. 3b). In the VMH-AHN-coll connectome, we observed a pronounced increase in membrane potential during oHFS, indicative of enhanced excitatory responses, primarily in inhibitory PRT clusters 8 and 10. Interestingly, spontaneous firing activity decreased in a cluster-specific manner, while diffuse increases in spontaneous firing were mapped across the tSNE distribution of VMH-AHN-coll PRTs. Furthermore, the frequency of action potential firing within bursts significantly increased (Fig. 3c). Notably, at both synaptic ends of the VMH-coll pathway, inhibitory responses silencing spontaneously active neurons or inducing rebound burst firing shifted toward excitatory profiles (Fig. 3d). In the VMH-AHN-coll connectome, the decrease in spontaneous firing was accompanied by increased excitation, suggesting that inhibitory responses pulled neurons closer to the GABA reversal potential, thereby enhancing electrochemical driving forces for excitatory synaptic inputs (Fig. 3e). Moreover, oHFS induced persistent changes in both postsynaptic connectomes, establishing a stable circuit state that did not revert to baseline. This stable state was consistently observed across both connectomes, suggesting a shared mechanism underlying oHFS-induced plasticity (Fig. 3f,g).

**Fig. 3:**
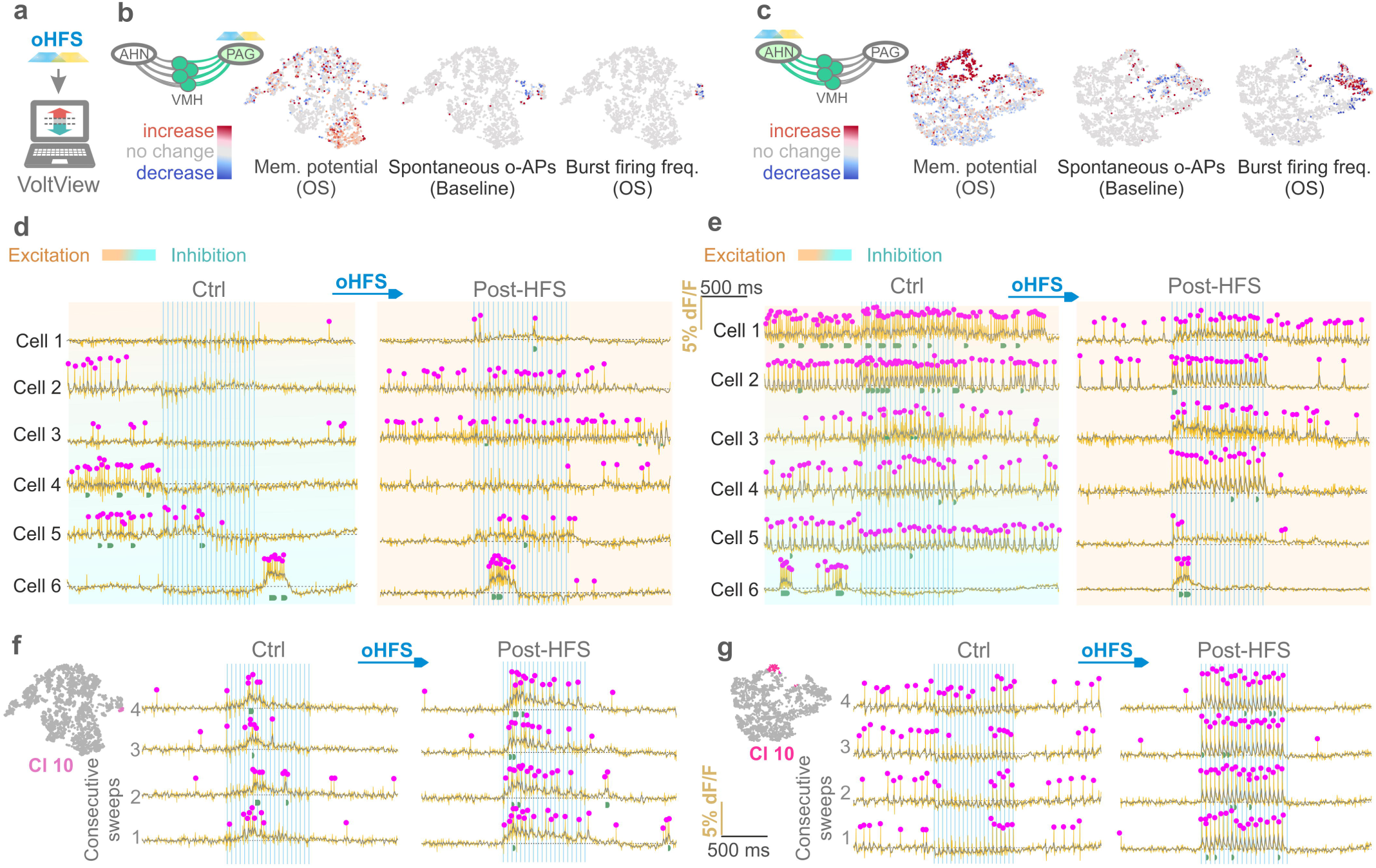
Circuit plasticity of the VMH collateral connectomes. All Voltron traces were reversed. **a,** Scheme of VoltView analysis to compare postsynaptic responses after oHFS. **b,** Scheme illustrating all-optical voltage imaging of the VMH-PAG-col connectome (left), t-SNE plots of increase (red) decrease (blue) or no change (gray) of membrane potential, spontaneous o-AP firing and firing frequency during bursts for each imaged neuron (right). **c,** Scheme illustrating all-optical voltage imaging of the VMH-AHN-col connectome (left), t-SNE plots of increase (red) decrease (blue) or no change (gray) of membrane potential, spontaneous o-AP firing and firing frequency during bursts for each imaged neuron (right). **d,** Example all-optical imaging from six PAG neurons (Neuron 1-6) of the VMH-PAG-col postsynaptic connectome before (Ctrl) and after oHFS (Post-oHFS), background color indicates the excitatory (bronze) or inhibitory (cyan) nature of the PRTs. **e,** Example all-optical imaging from six AHN neurons (Neuron 1-6) of the VMH-AHN-col postsynaptic connectome before (Ctrl) and after oHFS (Post-oHFS), background color indicates the excitatory (bronze) or inhibitory (cyan) nature of the PRTs. **f,** Example of a PAG neuron with bursting PRT from VMH-PAG-col cluster 1 before (Ctrl) and after oHFS (Post-oHFS) with 4 consecutive sweeps acquired through 1 minute showing a stable change of the PRT. **g,** Example of a PAG neuron with inhibitory PRT from VMH-AHN-col cluster 10 before (Ctrl) and after oHFS (Post-oHFS) with 4 consecutive sweeps acquired through 1 minute showing a stable excitatory change of the PRT.

### VMH collateral neurons drive defensive behaviors

Besides our finding of the quantitative bias of the VMH-coll pathway towards the AHN, both postsynaptic targets expressed synaptic plasticity and could establish a circuit state. In the VMH-PAG-coll connectome the strengthening of initially weak connections upon oHFS could indicate the existence of low release probability networks in the PAG, similar circuit properties have been reported earlier^28^, that need robust input to drive behavior. Therefore, we hypothesized that the VMH-PAG-coll pathway could evoke defensive behavior. To test our hypothesis, we performed *in vivo* optogenetics to activate the synaptic terminals of the VMH-PAG-coll axons. We labeled the VMH-PAG-coll axons with a dual viral labeling strategy by injecting retro-AAV-Cre into AHN and Cre-dependent AAV-DIO-ChR2-eYFP into the VMH (Fig. 4a) and we implanted optic fibers above the PAG. Remarkably, optogenetic activation of VMH-PAG-col terminals elicited pronounced immobility and freezing (Fig. 4b). By using 3 minutes ON/ OFF optogenetic activation periods we could observe heightened anxiety-like grooming and wall rearing between ON periods (Fig. 4b). Immobility increased across sessions within a day (Fig 4b), thus we tested how behavior would change by multiple activations on consecutive days. We found that immobility driven by the VMH-PAG-coll pathway progressively increased the time spent immobile across days (Fig 4f). Since biphasic behavior initiating with freezing and transitioning into jumping and escape was previously reported^10^ and the VMH-PAG-col pathway drove strong immobility we aimed to understand whether the VMH-collateral pathway promotes one or multiple different defensive strategies. We directly tested optogenetic activation of VMH-PAG-coll somas *in vivo.* To selectively label the VMH-coll somas we injected retro-AAV-Cre to the AHN, retro-AAV-Flpo to the PAG and Con/Fon-ChR2-eYFP into the VMH and implanted optic fiber above the VMH (Fig. 4c). Upon optogenetic activation of the VMH-coll somas, mice exerted jumps, escape and wall rearing during the 3 minutes ON periods and displayed heightened grooming between optogenetic stimulations (Fig. 4d). VMH-coll somatic activation induced only one type of defensive behavior. However, based on our all-optical measurements of synaptic plasticity in both VMH-coll connectomes and the induced behavior by the VMH-PAG-coll axons, we aim to understand whether repeated activation of VMH-coll somas would induce different potentiation on the two synaptic ends resulting in a transition between defensive behaviors. Upon repeated optogenetic activation of the VMH-coll somas across D1-D3 consecutive days, we observed decrease of jumps (Fig. 4g) and an escalation of escape with higher speed during ON periods (Fig. 4e), but without any change in the amount of immobility (Fig. 4f). We also tested whether three days of VMH-coll pathway activation induced an anxiety-like state. Both looming test and elevated plus maze showed significant difference from the control group (Fig 4h). Furthermore, we found that predator scent stress (PSS) with bobcat urine, that activates defensive immobility, induced early gene activation co-localizing with the VMH-coll neurons (Fig 4i). We asked whether the increased escape responses trained by the repeated activation of the VMH-coll pathway would change the innate immobility response activated by naturalistic olfactory predator cue. We found that mice exposed to bobcat urine after three days of VMH-coll optogenetic activation (Fig 4j) reversed defensive immobility to running escape (Fig. 4k). Moreover, mice displayed elevated running speed across all three days, indicating an induced brain state (Fig. 4k,l). Interestingly, immobility did not change across days compared to mice without opto-trained VMH-coll pathway that showed an increase in PSS-induced immobility across days (Fig. 4l). We mapped the c-FOS activation in the VMH of animals that were only exposed to bobcat urine and of those that were exposed to PSS after VMH-coll opto-training (Fig. 4m), and we found significantly stronger VMH activation in the latter group that was not restricted to the VMH-coll density core but affected other subregions of the VMH (Fig. 4n). These results revealed the role of VMH-coll pathway in the behavioral plasticity of innate defensive strategies.

**Fig. 4:**
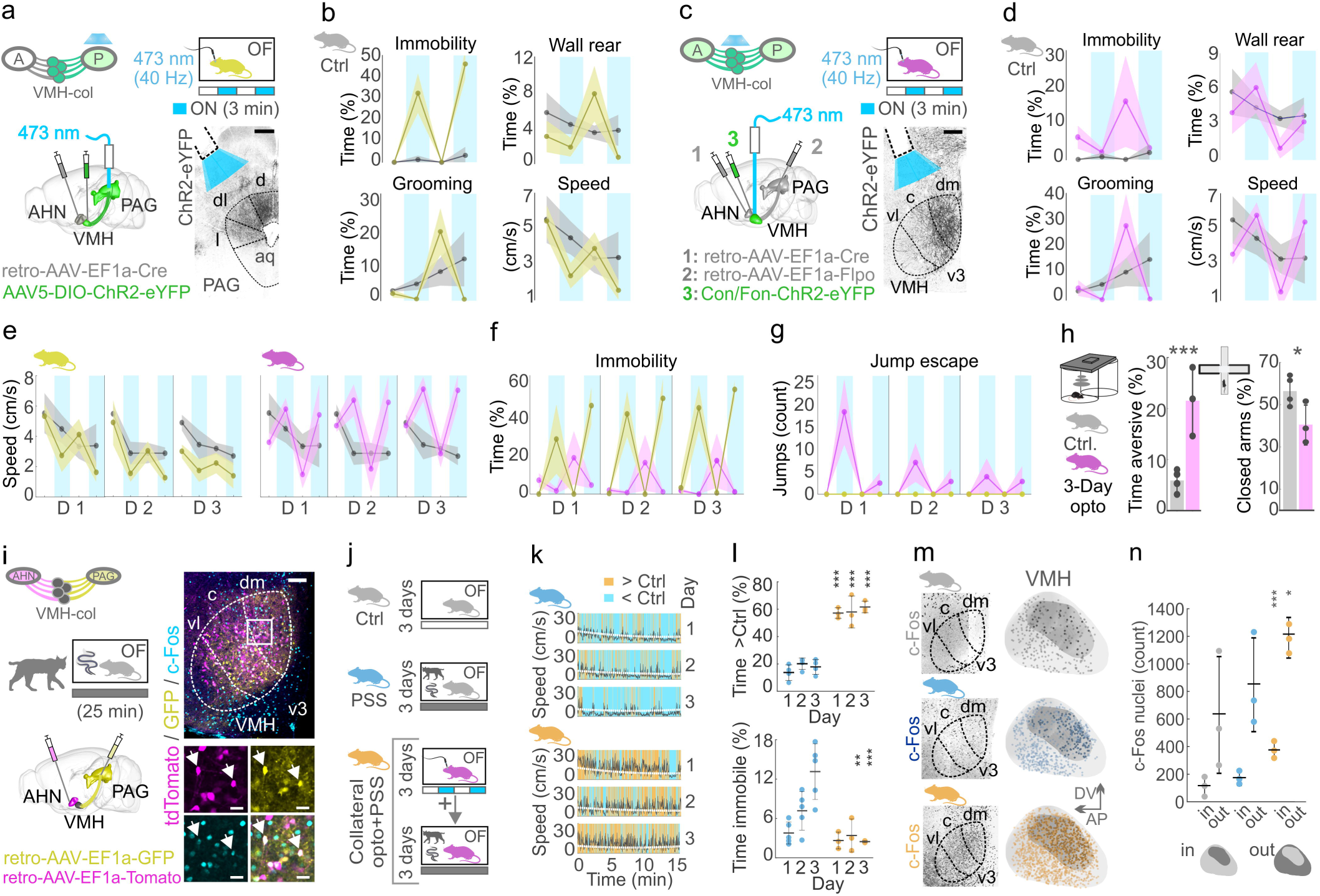
VMH collateral neurons drive defensive behaviors. **a,** Schematic of viral expression of ChR2 in the VMH-PAG-coll pathway and optical fiber placement above the PAG (left), alongside 3 min ON/OFF in vivo optogenetic activation of VMH-PAG-coll terminals in the OF (yellow group, N = 5, top right). Confocal image shows ChR2-eYFP expression with an implanted optic fiber above the PAG (right, scale bar: 500 µm). **b,** Scatter plots with sd± for immobility, wall rearing, grooming, and overall speed compared to control (gray, N = 4) during VMH-PAG-coll terminal activation. **c,** Schematic of viral expression of ChR2 in the VMH-coll somas and optic fiber placement above the VMH (left), alongside 3 min ON/OFF optogenetic activation of VMH-col somas in the OF (purple group, N = 3, top right). Confocal image shows ChR2-eYFP expression with an implanted optic fiber above the VMH (right, scale bar: 200 µm). **d,** Scatter plots with sd± for running, rearing, immobility, and grooming compared to control (gray) during VMH-coll soma activation. **e,** Scatter plots with sd± of speed in the yellow group across three consecutive days of VMH-PAG-col terminal activation compared to control (gray), and of VMH-coll soma activation in the purple group compared to control (gray). **f,** Scatter plots with sd± for immobility in VMH-PAG-coll-activated mice (yellow) compared to VMH-coll soma-activated mice (purple). **g,** Scatter plots with sd± for jump escapes comparing yellow and purple groups. **h,** Bar plot of total aversive behaviors during the looming test (left) and in the EPM (right) after three days of VMH-coll soma activation in the purple group compared to control (gray). **i,** Schematic of VMH-coll somatic anatomical labeling and PSS with bobcat urine (left). Confocal image shows VMH-PAG (yellow), VMH-AHN (magenta), and VMH-coll somas (white colocalization) with c-Fos immunostaining (cyan, scale bar: 100 µm). White rectangle indicates inset shown on separate channels below (scale bars: 20 µm). **j,** Schemes illustrating the control (gray), PSS (blue), and three-day VMH-coll soma activation followed by three consecutive days of PSS exposure (orange) groups compared in subsequent panels. **k,** Line plots of speed traces (black) during 15 min PSS in the blue (N = 4) and orange (N = 3) groups, with fitted control speed (white dashed line). Two-second binned points are colored orange if above control and blue if below control speed across three days (Day 1, 2, 3). **l,** Bar plots of time above control (top) and time spent immobile (bottom) during 15 min PSS in blue and orange groups across Days 1, 2, and 3. **m,** Confocal images of c-Fos immunostaining (left) with spatial map of c-Fos signal in the VMH of gray, blue, and orange groups. **n,** Bar plot quantifying c-Fos in the collateral volume (in) and outside of it (out) in the VMH of the gray, blue, and orange groups.

### *Esr1*-expressing VMH collateral neurons drive defensive immobility

Since we found that extreme activation of the VMH-coll pathway did not transition behavior from escape to immobility or vice versa, but the VMH-PAG-coll pathway induced strong immobility we hypothesized that different neuronal subpopulations could be responsible for driving the different defensive behaviors. We took advantage of the whole-brain RNA-transcriptome data recently released^29^ and the Allen *in situ hybridization* (ISH) database and found *Esr1* gene as a differentially expressed marker of the dm/cVMH that overlaps with the VMH-coll density core. Neurons expressing SF-1 are identified to be exclusively VMH neurons within the hypothalamus^9^ and have been shown to drive the defensive behaviors that we also observed^2,11,30^. Analyzing the RNA-transcriptome of SF-1 hypothalamic neurons confirmed the expression of the *Esr1* gene, even in multiple molecular clusters of the SF-1 neurons (Fig5. a). To directly test whether VMH-coll neurons are expressing *Esr1* we developed a PhP.eB virus that expresses nuclear blue fluorescent protein (BFP) in a Cre-dependent manner. We anatomically labeled the VMH-coll pathway with retro-AAVs injections to the PAG and to the AHN and we delivered the PHP.eB-DIO-BFP with a retro-orbital injection (Fig. 5b). Cre-dependent BFP expression labeled large population of Esr1+ neurons in the VMH (Fig. 5c). Colocalization of the VMH-coll neurons and the nuclear BFP confirmed the *Esr1* expression in the VMH-coll pathway (Fig. 5d). Next, we used Cre-dependent retro-AAV-tdTomato to label the *Esr1*+ collateral neurons from PAG (Fig. 5e). We found that approximately 90% of the *Esr1*+VMH-coll neurons resided in the anterior dmVMH and not in the posterior portion of the VMH-coll density core (Fig. 5f,g). To probe the postsynaptic connectivity of the Esr1+VMH-coll neurons we systemically expressed Voltron-ST by retro-orbital injection of PhP.eB-Voltron-ST, and ChR2 by injecting retro-AAV-Flpo to the AHN and retro-AAV-Con/Fon-ChR2-eYFP to the PAG of *Esr1*-Cre mice (Fig 5h). This injection strategy also revealed the strong axonal coverage both in the *Esr1*+VMH-AHN-coll and *Esr1*+VMH-PAG-coll pathways (Fig. 5i). We employed all-optical voltage imaging of both *Esr1*-expressing VMH-coll pathways and similarly to imaging the wild-type VMH-coll connectomes, we tile-imaged the entire AHN and VMH on serial brain sections *ex vivo* (Fig. 5j). We used cluster load analysis to classify the PRTs captured in both *Esr1*+ VMH-coll connectomes (Fig. 5k). Comparison of 3931 neurons from wild-type and 9584 neurons from *Esr1*-expressing VMH-AHN-coll connectome showed two-third of the responses proportionally (Fig. 5l). The *Esr1*+ connectome qualitatively differed in four clusters, interestingly mainly the ones with inhibitory PRTs (Fig. 5m). The same type of comparison of 3212 neurons from wild-type and 5424 neurons from *Esr1*-expressing VMH-PAG-coll side (Fig. 5n) showed very high degree of similarity on both proportional quantitative and qualitative levels (Fig. 5o), suggesting that the *Esr1*+ VMH-coll pathways could represent the PAG driving pathway that could potentially induce defensive immobility. To probe the behavioral role of the *Esr1*+VMH-coll pathway, we used identical ChR2 expression strategy that we used for all-optical voltage imaging and implanted optic fibers above the VMH of *Esr1*-Cre mice (Fig. 5p). Remarkably, optogenetic activation of the *Esr1*+VMH-coll somas in an open field induced strong immobility (Fig. 5q,r) and similarly to VMH-PAG-coll terminal activation, grooming increased between optogenetic activations. Surprisingly, the activation of the *Esr1*+VMH-coll somas also induced flattening behavior that has not yet been associated with the VMH (Fig. 5s,t). Both freezing and flattening showed strong significance compared to control mice (Fig. 5u). These results unveiled the role of the newly identified Esr1+VMH-coll pathway in promoting innate defensive immobility and flattening.

**Fig. 5:**
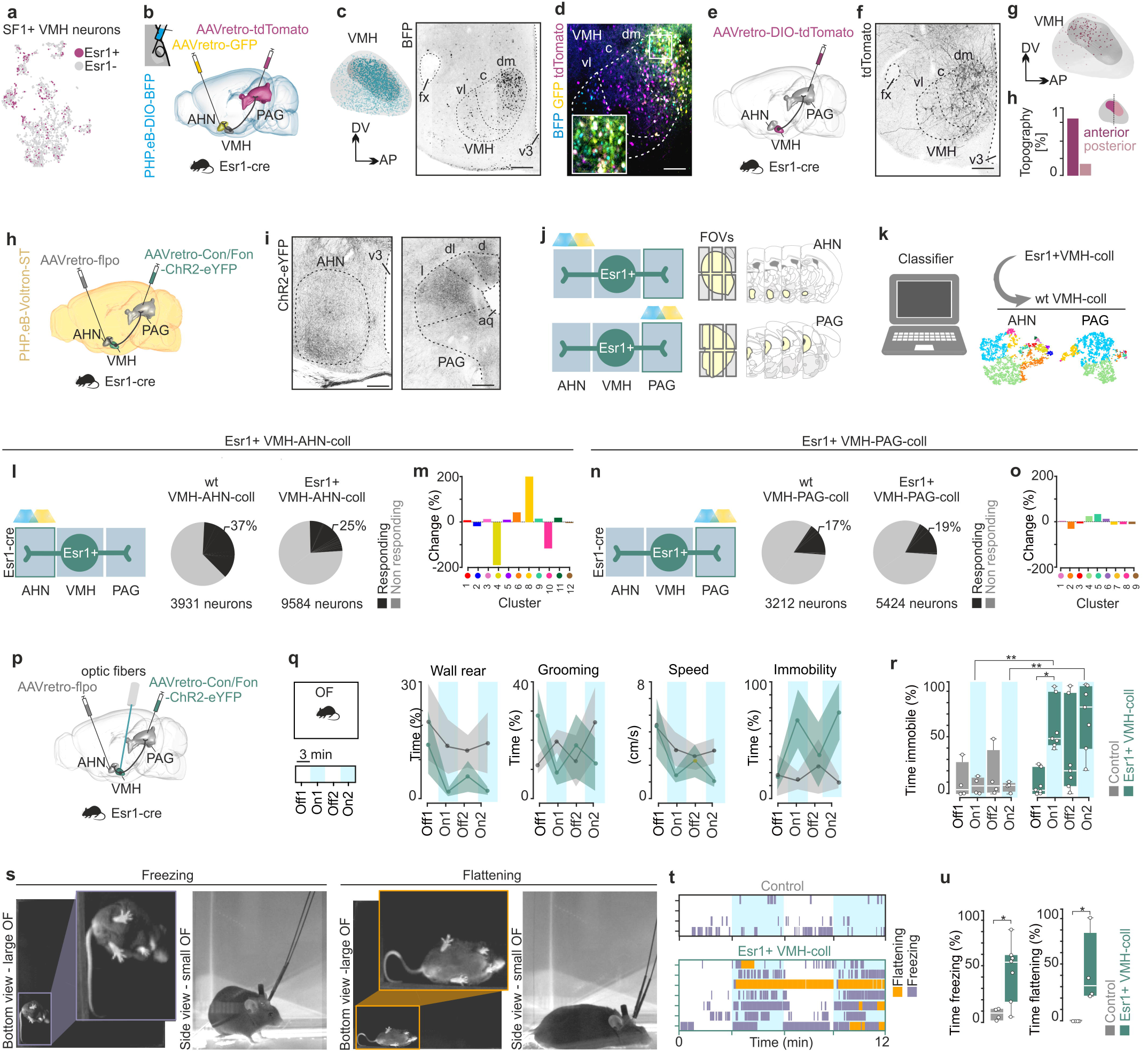
*Esr1*-expressing VMH collateral neurons drive defensive immobility. a, tSNE plot of single-cell RNA-transcriptome of SF1-expressing VMH neurons with overlayed *Esr1* expression (hot pink). **b,** Scheme of anatomical labeling of VMH-coll pathways with retro-AAVs from PAG and AHN combined with retro-orbital virus injection for systemic labeling of *Esr1*+ neurons. **c,** Spatial VMH map of BFP signal identifying *Esr1*+ neurons in the VMH (blue, left), confocal image of BFP expression in the VMH at bregma −1.35 (right). **d,** Confocal image showing the colocalization of collateral neurons and Cre-dependent BFP in Esr1-Cre mouse **e,** Scheme of retrograde Cre-dependent viral labeling of *Esr1*+ VMH-coll neurons in *Esr1*-Cre mouse. **f,** Confocal image of Esr1+ VMH-coll neurons. **g,** Spatial mapping of *Esr1*+ VMH-coll neurons located in the anterior portion of the VMH-coll volume (top), bar plot of comparing spatial topography of *Esr1*+ VMH-coll neurons. **h,** Scheme of viral labeling strategy to express ChR2 in the *Esr1*+ VMH-coll neurons and Cre-independent systemic Voltron-ST for large anatomical coverage in *Esr1*-Cre mice. **i,** Confocal images of ChR2-expressing axonal terminals in the AHN (left) and in the PAG (right). **j,** Scheme of all-optical voltage imaging of Esr1+VMH-AHN-coll and Esr1+VMH-PAG-coll postsynaptic connectomes with tile-imaging to cover the entire AHN and PAG on serial coronal brain slices *ex vivo*. **k,** Illustration of the classifier analysis that compared all-optical recorded *Esr1*+VMH-coll PRTs and wild-type VMH-coll all-optical-recorded PRTs. **l,** Scheme illustrating VMH-AHN-coll all-optical connectome imaging (left), pie chart of classified PRTs in 3931 neurons of the VMH-AHN-coll (left) and 9584 neurons of the *Esr1*+VMH-AHN-coll (right) connectomes (right). **m,** Bars of weighted proportional difference of cluster load analysis comparing *Esr1*+VMH-AHN-coll and VMH-AHN-coll connectomes. **n,** Scheme illustrating VMH-PAG-coll all-optical connectome imaging (left), pie chart of classified PRTs in 3212 neurons of the VMH-PAG-coll (left) and 5424 neurons of the *Esr1*+VMH-PAG-coll (right) connectomes (right). **o,** Bars of weighted proportional difference of cluster load analysis comparing *Esr1*+VMH-PAG-coll and VMH-PAG-coll connectomes. **p,** Scheme of viral labeling strategy to express ChR2 in the *Esr1*+ VMH-coll neurons and implantation of optic fiber above the VMH in an *Esr1*-Cre mouse. **q,** Scheme of open field (OF) experiment with 3 min ON/OFF epochs to optogenetically activate *Esr1*+VMH-coll pathway *in vivo* (left), scatter plots with sd± of behaviors during 3 min ON/OFF optogenetic experiment quantifying wall rearing, grooming, speed and immobility (right). **r,** Box plot of defensive immobility during 3 min ON/OFF optogenetic experiment in control (gray) and *Esr1*+VMH-coll activation (green). **s,** Example video frames of the observed freezing (left, purple) and flattening behavior (right, orange). **t,** plot of freezing and flattening during the optogenetic activation of *Esr1*+VMH-coll pathway. **u,** Box plots quantifying the time spent freezing (left) and flattening (right) compared to control (gray).

### Peptide neuromodulation of the *Esr1*+ VMH collateral connectomes

To understand the mechanisms that induce circuit plasticity in the VMH-coll connectomes we aimed to investigate the interplay of fast synaptic neurotransmission and peptide neuromodulation that shapes synaptic connectivity. We used the open-source whole-brain RNA-transcriptome data to analyze the expression of neuropeptides in the SF1-expressing VMH neurons and found that *Chga* (Chromogranin A), *Adcyap1* (Adenylate Cyclase Activating Polypeptide 1), *Pdyn* (Prodynorphin), *Tac1* (Tachykinin Precursor 1), and *Pomc* (Pro-opiomelanocortin) had the highest expression (Fig. 6a). All these neuropeptides mapped uniquely to the tSNE of SF1-expressing molecular clusters and overlapped with *Esr1* expression (Fig. 6b). We aimed to understand the role of peptide neuromodulation and investigated how PACAP influences the Esr1+VMH-coll postsynaptic connectomes. PACAP had been reported to be expressed primarily by excitatory glutamatergic neurons^31^ potentially modulating glutamatergic transmission. We performed all-optical voltage imaging of the postsynaptic connectome *ex vivo* and applied neuropeptide pharmacology to probe neuromodulation of the Esr1+VMH-coll connectomes. We found overall excitatory effect in the AHN indicated by the increase in spontaneous firing frequency and increased response efficacy of AP firing during OS (Fig. 6c). Interestingly the Esr1+VMH-PAG-coll connectome had the opposing response with decrease in the spontaneous firing indicative of hyperpolarization of PAG neurons (Fig. 6c). These findings revealed that neuropeptide modulation can exert different effects in collateral connectomes. We found high levels of dynorphin in the SF1-expressing VMH population, therefore we aimed to probe its neuromodulatory effects on the Esr1+VMH-coll connectomes, as dynorphin has been reported to mediate stress-induced dysphoria via the KOR system^22,23^. We performed all-optical voltage imaging of the Esr1+VMH-coll connectomes *ex vivo* and bat-applied neuropeptides to probe neuromodulation. We found that dynorphin exerted depolarizing effect on the AHN synaptic end of the Esr1+VMH-coll pathway, increased spontaneous firing and AP firing during OS (Fig. 6d). Interestingly, we observed that besides robust potentiation of excitatory connections, inhibitory PRTs have been transitioning to excitatory, a similar effect we observed after oHFS of the VMH-coll connectomes (Fig. 6e). We found surprisingly similar dynorphin effect in the Esr1+VMH-PAG-coll connectome with increased AP firing that was maintained even after OS (Fig. 6f,g). Our results suggested that because of the similar effects of dynorphin on the PRT changes of both synaptic ends to those we recorded *ex vivo* after oHFS, mimicking high level of pathway activation, we aimed to understand the role of dynorphin neuromodulation of the Esr1+VMH-coll pathway *in vivo*. We expressed ChR2 in the Esr1+VMH-coll pathway by injecting retro-AAV-Flpo to the AHN and retro-AAV-Con/Fon-ChR2-eYFP to the PAG of *Esr1*-Cre mice and implanted optic fiber above the VMH of *Esr1*-Cre mice (Fig. 6h). To occlude the neuromodulatory effect of dynorphin, we i.p.-applied Aticaprant, antagonist of KOR system, 24 hours before pathway stimulation (Fig. 6i). VMH-coll pathway activation at both synaptic ends, and with both terminal and somatic stimulations resulted in an increase of the expression of the elicited behaviors across optogenetic activation sessions (Fig. 4,5). Remarkably, optogenetic activation of the Esr1+VMH-coll in the presence of KOR antagonist exerted less pronounced potentiation of the evoked defensive immobility (Fig. 6j,k). Already within one day, the change of speed between the first and the second 3-min long optogenetic activation was significantly higher in the Aticaprant-treated mice (Fig. 6l). Systemic antagonism of the KOR signaling not only resulted in less pronounced immobility, but also significantly delayed the onset of immobility (Fig. 6m). Our findings revealed that dynorphin neuromodulation of the Esr1+VMH-coll pathway is necessary for efficient behavioral plasticity of innate defensive immobility.

**Fig. 6:**
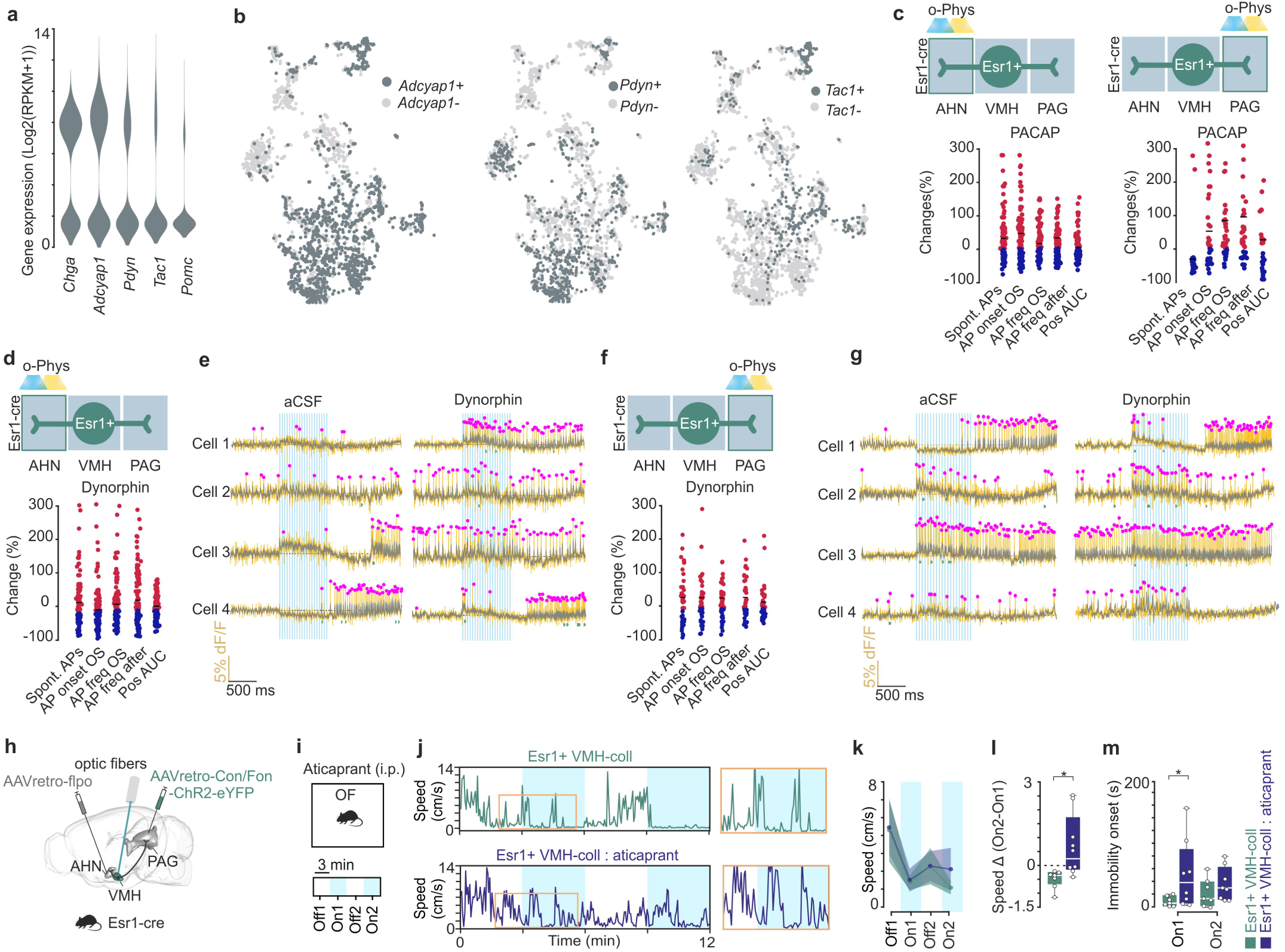
Peptide neuromodulation of the *Esr1*+ VMH collateral connectomes. Voltron traces were reversed **a,** Violin plots displaying the gene expression levels of neuropeptides within the SF1-expressing VMH neurons, *Chga* (Chromogranin A), *Adcyap1* (Adenylate Cyclase Activating Polypeptide 1), *Pdyn* (Prodynorphin), *Tac1* (Tachykinin Precursor 1), and *Pomc* (Pro-opiomelanocortin) **b,** t-SNE plots illustrating the distribution of *Adcyap1*, *Tac1*, and *Pdyn* expression in SF1-expressing VMH neurons; dark gray points indicate positive, light gray points indicate negative neurons. **c,** Scatter plots expressing changes (increase red, decrease blue) of o-phys parameters following the bath-application of PACAP (1-38) in AHN and PAG. **d,** Scatter plots expressing changes (increase red, decrease blue) of o-phys parameters following the bath-application of dynorphin **e,** Example traces before (left) and following dynorphin wash on (right), from four different PAG neurons **f,** Scatter plots expressing changes (increase red, decrease blue) of o-phys parameters following the bath-application of dynorphin in the AHN. **g,** Example traces before (left) and following dynorphin wash on (right), from four different AHN neurons. **h,** Scheme of viral labeling strategy to express ChR2 in the *Esr1*+VMH-coll neurons and implantation of optic fiber above the VMH in an *Esr1*-Cre mouse. **i,** Scheme of the experimental strategy to evaluate dynorphin-mediated *Esr1*+VMH-coll pathway neuromodulation *in vivo*. Aticaprant (10mg/Kg i.p.) was administered 24 hours before opto-open field (OF) experiment. **j,** Representative speed traces for *Esr1*+VMH-coll (green) and *Esr1*+VMH-coll + Aticaprant (purple). **k**, Scatter plots with sd± of speed during 3 min ON/OFF optogenetic experiment. **l**, Speed Δ On2-On1 was analysed (Student’s t-test, *p<0.05). **m**, Box plot of defensive immobility onset (sec) during the optogenetic activation of *Esr1*+VMH-coll with (purple) or without (green) Aticaprant treatment (two-way ANOVA, Šidák’s multiple comparison test, *p<0.05).

## Discussion

The findings presented in this study significantly advance our understanding of the ventromedial hypothalamus (VMH) and its role in modulating defensive behaviors. The study highlights the complex synaptic architecture and functional diversity of the VMH-collateral (VMH-coll) pathway, with distinct connectivity patterns toward the anterior hypothalamic nucleus (AHN) and periaqueductal gray (PAG). The differential engagement of these postsynaptic targets underscores the role of the pathway in orchestrating diverse defensive responses, from freezing to escape behaviors.

The research also sheds light on the anatomical and functional specificity of Esr1-expressing VMH-coll neurons. These neurons preferentially drive immobility and flattening behaviors, contrasting with the escape responses elicited by broader VMH-coll activation. The use of advanced optogenetic and imaging techniques to map these pathways provides a nuanced understanding of how distinct neuronal populations within the VMH contribute to behavior. Moreover, the study demonstrates that systemic antagonism of the KOR system diminishes behavioral plasticity, underscoring the importance of dynorphin in maintaining the flexibility of defensive responses.

Future research could aim to dissect how the VMH integrates multisensory inputs to prioritize behavioral responses could provide insights into how innate survival strategies are shaped by experience. Additionally, exploring the role of other neuromodulators, such as PACAP or SP in fine-tuning VMH-coll circuitry could reveal complementary mechanisms underlying behavioral adaptability. Finally, extending these findings to pathological conditions, such as anxiety disorders or post-traumatic stress disorder (PTSD), may pave the way for targeted therapies that modulate VMH-coll pathways to restore adaptive behavioral responses. This study provides a robust framework for understanding the role of collateral VMH in behavioral plasticity, offering exciting avenues for future exploration in both basic and translational neuroscience.

## Methods & Protocols

### Animals

All procedures and experiments on animals were performed according to the guidelines of the Stockholm Municipal Committee for animal experiments and the Karolinska Institutet in Sweden (approval no 17073-2023). Experiments were conducted using adult 3-6 months male and female mice, wild type C57BL/6J (Charles River Laboratories) or the transgenic mouse line: Esr1-cre (B6N.129S6(Cg)-Esr1tm1.1(cre)And/J; Jackson Laboratory Stock, 017911). All transgenic mice used in experiments were heterozygous for the transgenes. Mice were group housed, up to five per cage, in a temperature (23°C) and humidity (55%) controlled environment in standard cages on a 12:12hr light/dark cycle with *ad libitum* access to food and water.

### Viral constructs

All purified and concentrated adeno-associated viruses (AAV) were purchased from Addgene.

### Anatomy and histology

pAAV-CAG-GFP (Addgene, Cat. No.: 37825-AAVrg)

pAAV-CAG-tdTomato (Addgene, Cat. No.: 59462-AAVrg)

### In vivo experiments

AAVrg-EF1a-Cre (Addgene, Cat. No.: 55636-AAVrg)

pAAV-EF1a-doublefloxed-hChR2(H134R)-EYFP-WPRE-HGHpA (Addgene, Cat. No.: 20298-AAV5)

pAAV-EF1a-Flpo (Addgene, Cat. No.: 55637-AAVrg)

hSyn-Con/Fon-hChR2(H134R)-EYFP (Addgene, Cat. No.: 55645-AAVrg)

### Custom made viral construct for anatomy and voltage imaging

ssAAV-PHP.eB/2-hSyn1-chI-Voltron-ST-WPRE-bGHp(A) (VVF Zurich, Cat. No.: v827-PHP.eB) ssAAV-PHP.eB/2-hSyn1-chI-dlox-mTagBFP2_NLS(rev)-dlox-WPRE-SV40p(A) (VVF Zurich, Cat.No.: v883-PHP.eB)

### Viral injections

#### General procedure

Mice were anesthetized with isoflurane (2%) and placed into a stereotaxic frame (Harvard Apparatus, Holliston, MA). Before the first incision the analgesic Buprenorphine (0.1 mg/kg) and local analgesic Xylocain/Lidocain (4 mg/kg) was administered subcutaneously. The body temperature of the mice was maintained at 36 °C with a feedback-controlled heating pad. For viral injections a micropipette attached on a Quintessential Stereotaxic Injector (Stoelting, Wood Dale, IL) was used. Injections were done with the speed of 50 nl/minute. The injection pipette was held in place for 5 min after the injection before being slowly (100 µm/s) retracted from the brain. The analgesics Carprofen (5 mg/kg) was given at the end of the surgery, followed by a second dose 18-24h after the surgery.

#### Labeling strategies

For anatomical characterization and electrophysiological recordings of the VMH-PAG pathway in C57BL/6J mice, 0.3μl pAAV-CAG-GFP or pAAV-hSyn-hChR2(H134R)-EYFP was unilaterally injected into the VMH (coordinates: AP −1.45 mm, ML 0.25 mm, DV −5.25 mm). Targeting of the PAG was achieved by 1 (coordinates: AP −4.1 mm, ML 0.2 mm, V −1.6 mm) or 2 (coordinates: AP −3.9 mm and AP −4.3 mm, ML 0.2 mm, V −1.6 mm) unilateral injections of 0.3μl of 1:1 mixture of pENN-AAV-hSyn-Cre-WPRE-hGH and pAAV-hsyn-flex-Voltron-ST.

### Histology

#### General procedure

Mice were deeply anaesthetized with Na-pentobarbital (60mg/kg) and transcardially perfused with 0.1M PBS followed by 4% paraformaldehyde in PBS 0.1M. Brains were removed and post-fixed in 4% paraformaldehyde in PBS 0.1M overnight at 4°C and then washed and stored in 0.1M PBS. Coronal, 50 μm slices were cut using a vibratome (Leica VT1000, Leica Microsystems, Nussloch GmbH, Germany). The sections were washed in 0.1M PB and mounted on glass slides (Superfrost Plus, Thermo Scientific) and coverslip-covered(Thermo Scientific) using glycerol: 1x PBS (50:50).

#### Histology of biocytin filled neurons

250 μm thick brain slices containing biocytin-filled neurons and voltron-JF585 labeling were post-fixed in 4% paraformaldehyde in phosphate-buffer (PB, 0.1M, pH 7.8) at 4 °C overnight. Slices were repeatedly washed in PB and cleared using CUBIC protocol^19^. First “CUBIC reagent 1” was used (25 wt% urea, 25 wt% N,N,N’,N’-tetrakis(2-hydroxypropyl) ethylenediamine and 15 wt% polyethylene glycol mono-p-isooctylphenyl ether/Triton X-100) for 1 day at 4 °C. After repeated washes in PB, biocytin was visualized using Alexa Fluor 633-conjugated streptavidin (1:1000, RT 3 hours). For NeuN staining primary antibody (Millipore, MAB377,1:1000) was incubated overnight at 4 °C and after repeated washing with PB, 2nd antibody (Jackson, Cy5, Code:715-175-151) was incubated for 3h RT. Slices were then re-washed in PB and submerged in “CUBIC reagent 2” (50 wt% sucrose, 25 wt% urea, 10 wt% 2,20,20’-nitrilotriethanol and 0.1% v/v% Triton X-100) for further clearing. Slices were mounted on Superfrost glass (Thermo Scientific) using CUBIC2 solution and covered with 1.5 mm cover glasses.

#### Confocal imaging

All confocal images were taken using a Zeiss 880 confocal microscope. CUBIC cleared sections after slice electrophysiology and biocytin or NeuN staining were acquired as z-stacks using a Plan-Apochromat 20x/0.8 M27 objective (imaging settings: frame size 1024×1024, pinhole 1AU, Bit depth 16-bit, speed 6, averaging 4). For viral expression overview of coronal cut VMH or PAG sections were acquired with a Plan-Apochromat 20x/0.8 M27 objective (imaging settings: frame size 1024×1024, pinhole 1AU, Bit depth 16-bit, speed 7, averaging 2). For ISH images oil-immersion 63x/1.0 objective was used (imaging settings: frame size 1024×1024, pinhole 1AU, Bit depth 16-bit, speed 6, averaging 4) Processing of images was either done in ImageJ (NIH, USA) or Imaris 7.4.2 (Oxford Instruments, UK).

### Brain slice preparation *ex vivo*

First, mice were anesthetized with intraperitoneal injection of 50 µl Na-pentobarbital (60 mg/kg) and transcardially perfused with 4-8 °C cutting solution, containing (in mM): 40 NaCl, 2.5 KCl, 1.25 NaH2PO4, 26 NaHCO3, 20 glucose, 37.5 sucrose, 20 HEPES, 46.5 NMDG, 46.5 HCl, 1 L-ascorbic acid, 0.5 CaCl2, 5 MgCl2. Next, brain was carefully removed and 250 μm thick coronal slices were cut with a vibratome (VT1200S, Leica, Germany) in the same 4-8 °C cutting solution. Next, slices were incubated in cutting solution at 34°C for 13 minutes, and kept until recording at room temperature in artificial cerebrospinal fluid (aCSF) solution containing (in mM): 124 NaCl, 2.5 KCl, 1.25 NaH2PO4, 26 NaHCO3, 20 glucose, 2 CaCl2, 1 MgCl2. For Voltron-imaging, slices were incubated at room temperature in Janelia-Fluor 585 HaloTag (JF-dyes, Janelia) ligands. JF-dyes were dissolved in DMSO to a stock of 1 µM and further diluted to 50 nM in aCSF before use. All solutions were oxigenated with carbogen (95% O2, 5% CO2). All constituents were from Sigma-Aldrich.

### Voltage imaging

After 4-5 weeks of Voltron-ST expression, mice were sacrificed and ex vivo brain slices of 250 µm were prepared. After ∼30 min of incubation in 50 nM Janelia-Fluor 585 dye dissolved in aCSF, slices were transferred to the recording chamber of the electrophysiology setup. For the imaging we used a digital sCMOS camera (Orca Fusion-BT, Hamamatsu, Japan), frame triggers were sent to the camera with a Arduino Micro microcontroller (Arduino Uno) with 600 Hz. For all-optical imaging we used a dual-band excitation filter (ZET488/594, Chroma) to excite the JF-585 and deliver 473 nm light for optogenetic stimulation. The 585 nm light excitation intensity was ∼10 mW/mm^2^ and 473 nm light intesity was ∼2.5 mW/ mm^2^ at the slice plane and was delivered by Spectra X (Lumencore) LED light source. JF-585 fluorescent emission was collected with a 20X 1.0 NA water immersion objective (XLUMPLFLN20XW Plan Fluorit, Olympus). Emitted light was separated from the excitation light with a band-pass emmission filter (ET645/75, Chroma) and with a dichroic mirror (T612lprx, Chroma). Magnification was decreased with U-ECA magnification changer to 0.5x. To acquire videos we used the free LiveImage software triggered by the Arduino to synchronize acquisition with the frame triggers.

### In vivo optogenetics

Mice were bilaterally implanted with optical fibers targeting the VMH (coordinates: −1.45 mm AP, 1.25 mm ML from bregma, and 4.8 mm depth at a 10° angle from the dura). The implanted optical fibers were purchased from RWD Life Science (R-FOC-BL200C-22NA, 3 mm for LHb, 5 mm for LHA). Mice were connected via a splitter branching patch cord (SBP(2)_200/220/900-0.22_1m_FCM-2xMF1.25, Doric Lenses) to their implanted optical fibers using a split sleeve (ADAL1-5, Thorlabs). The splitter branching patch cord was connected to a laser (MLL-III-447-200mW laser) via a fiber-optic rotary joint (FRJ_1×1_FC-FC, Doric Lenses) to prevent cable twisting during animal’s movement. After each testing session, the animals were uncoupled from the splitter branching patch cord and returned to their home cages. The frequency and duration of photostimulation were controlled using a custom-written Arduino script (Arduino IDE), managed through Bonsai software v2.6.3. For optogenetic stimulation, the power was measured in continuous light at 8 mW, measured at the tip of the splitter branching patch cord cable before each experiment using an optical power and energy meter with a photodiode power sensor (Thorlabs). The stimulation was carried in continuous light of 40Hz (5ms). Animals with undetectable viral expression in the target region or incorrect optical fiber placement were excluded from the analysis.

### Opto-open field test

Mice were handled for three consecutive days prior to behavioral experiments and acclimated to the behavioral room for 1 hour before testing. Behavior was recorded using a CCD camera (20 Hz frame rate) with an infrared filter, interfaced with Bonsai software v2.6.3. Only red and infrared light were used during all experiments. Mice were habituated to the laser cables for 3 days before testing. Mice were placed in a custom-made open field (30 x 30 cm) made of black plexiglass for 15 minutes. The arena was positioned on a transparent plexiglass surface to allow the camera to record behavior from below at 20 fps. Mouse performance was evaluated during alternating epochs of laser stimulation, beginning with a 3-minute off period, followed by 3-minute stimulation epochs (continuous light, 40 Hz, 5 ms pulse, 447 nm laser). Before testing a new mouse, the maze was cleaned with 70% ethanol. Body point pose estimation in open field test videos was conducted using DeepLabCut (DLC). Six virtual markers were placed on key body parts (nose, tail base, paws) across 375 sampled frames from 15 videos. A residual neural network (ResNet-50) was trained to track these markers. Speed was calculated using tail base marker. Immobility, flattening, grooming and rearing behaviors were scored manually. Data were presented in box plots, with individual data points overlaid to identify animals. Aticaprant (HY-101718, MedChemExpress, formerly LY-2456302) was dissolved 500µM in MilliQ water and titrated to pH 5. Aticaprant 5µM in saline solution was then prepared fresh prior to use. Aticaprant 10 mg/kg was injected intraperitoneally 24 hours before opto-open field test.

### Predator scent stress

Bobcat urine (PredatorPee® Bobcat Pee Brand Bobcat Urine) was used in the Smell group. 1 - 1,5 ml were used for each exposure. Mice were handled for three consecutive days before the behavioural experiments. Animal behaviour was recorded with infrared illumination, invisible to the mice. Excluding the open field (OF), all tests were performed on the same days as a behavioural assessment of the animals. They were performed in an increasing level of aversiveness: animals first completed the elevated plus maze (EPM) test, followed by the acoustic startle response test (ASRT), and finally the looming stimulus test (LST).

#### Open field test

Mice were placed in an OF box (45×45 cm) for 15 min. The behavioural arena was placed under the camera and the animal was recorded from above. There was no compulsive behaviour forced on the animal. A perforated petri dish (100 mm x 15 mm) containing a small petri dish (30 mm x 15 mm) inside, fixed in a corner of the box, was used for placing the bobcat urine in the smell group. For the *Ctrl* group, the small petri dish was empty. The same procedure was repeated for 4 consecutive days for repeated exposure, although there were changes in the context, such as differences in the light or the appearance of the box, to prevent contextual fear. Drugs were administered intraperitoneally (IP) 25 min before testing. The animal was recorded by a video tracking system (EthoVisionXT17). Later, its behaviour was analysed using DeepLabCut (DLC) and MATLAB. Animals repeated the Open Field test under the same conditions one last time, 20min before perfusing.

#### Elevated plus maze

Mice underwent an EPM assay (77 x 77 cm) as previously described ^32^. In brief, mice were placed at the junction of the four arms of the maze, facing an open arm, and left there for 15 min. This test is a widely used behavioural assay in rodents to evaluate the anti-anxiety effects of pharmacological agents ^32,33^. The animal behaviour was both recorded and analysed by a video-tracking system, EthoVision XT 17. The data obtained was analysed and plotted using MATLAB.

#### Acoustic startle response test

Mice were placed into a sound-isolated TSE Multi Conditioning System (MCS) – Fear Conditioning box. In the ASRT, as detailed in previous studies (53), animals were first left to habituate for 5 min. Next, they were exposed to 16 tones (5 seconds, 70 dB, 10 kHz) leaving an interval between tones randomly chosen (Fig. 4A). The animal behaviour was tracked by the local software MCS FCS – SQ – SM. The resulting data was analysed and plotted using MATLAB.

#### Looming stimulus test

Mice performed an LST, as previously described ^34^, in a custom-designed 8-shaped field with black matt walls to prevent reflection. The field resulted from merging two round arenas (30 cm diameter) with an opening between them to allow free exploration. The arena’s ceiling held a monitor emitting dim lighting from its grey screen simulating an approaching of aerial predators, which induce defensive reactions ^35,36^. The animal behaviour was recorded with a camera located below the arena, which was placed on a transparent plexiglass. The looming stimuli were triggered manually once the animal was in the centre of one of the arenas. In essence, the stimulus was repeated 15 times with increasing diameter (from 2 to 20 degrees of visual angle) during the first 250 seconds, and then it remained stable at 20 degrees for the remaining 250 ms. The next stimuli were triggered by the experimenter after, at least, 1 min from the previous one. The animal behaviour was recorded by a video tracking system (Bonsai), for later being analysed using DLC. The data obtained was analysed and plotted using MATLAB.

### Data analysis

#### VoltView analysis

The on-site and detailed analyis was written in MATLAB (MathWorks) using custom-written scripts. In major steps, the recorded videos from ‘.CXD’ files were imported to MATLAB using bioformats 6.11 package (OME - https://www.openmicroscopy.org/). Multiple sweeps were recorded in the same video in series. Putative lost frames were found and filled up. Camera artifact frames – marking the end of sweeps – were removed. Frames of the baseline before the optogenetic stimulation was averaged, frame average was local-equalized and contrast-enhanced specifically optimized with Voltron-JF585 imaging. These pre-processed images were used for the Cellpose segmentation of somatic ROIs. After the detection of ROI contours we designated the pixels of concentrical ROIs (Inner and Outer) around the somatic ROI. The number of pixels of Inner and Outer were scaled to be within ±5% of the somatic ROI surface so that signal-to-noise can be compared. We extracted the optical physiology (o-phys) traces from all somatic, Inner and Outer ROIs. We calculated the dF/F from the o-pys traces and used exponential fitting to correct for bleaching. Basic features, o-AP peaks, subthreshold kinetics (o-Sub) and Burst activity was detected first. O-AP detection used the noise of the baseline for each ROI to calculate a cutoff threshod at 2.5-times the std – this may differ based on the illumination light intensity of voltage imaging as the signal-to-noise ratio is affected by that. We used polynomial fitting for straightening the dF/F traces before the o-AP detection (which we also validated by data from simultaneous whole-cell patch-clamp recordings). O-sub was the moving average of the 600 Hz o-phys trace by default. As wider o-APs – based on o-AP-half-width measurement - needed different moving average, we scaled it accordingly for o-Sub detection. Burst detection used a slower moving average with averaging 100 points (o-Sub-slow) against the o-Sub, where the o-Sub went above the o-Sub-slow, the periods were considered as putative burst periods. Next, o-AP detection was overlayed and compared to putative burst periods and if multiple o-APs were on these fast depolarizations, they were considered to be bursts (this was optimized and validated with simultaneous whole-cell patch-clamp and all-optical voltage imaging as in Extended Data Fig 3c-e). Using the basic features (o-AP peaks, o-Sub, Burst) we compared the 3 o-phys traces of the ROIs (somatic, Inner, Outer) with the amplitude of o-APs, the amplitude of o-Sub and the length and number of bursts across the traces. If the amplitudes and burst length were decreasing going from the somatic ROI towards the Inner and towards the Outer zones, the ROI was comfirmed to be the source of the response. In case if the signal parameters became stronger from the soma towards the Outer zone, the ROI was discarded. Traces were denoised using wavelet transformation or with multiscale local 1-D polynomial transformation. Optical blue stimulation artifacts were removed. As the Voltron-imaging with our analysis could detect overall depolarization and hyperpolarization compared to baseline level during Op, oEPSPs and o-IPSPs were detected using the o-Sub thresholded with ±5 x sd of the baseline and searching changes in the first derivative which represent inreases of o-EPSPs or decreases of o-IPSPs. All the calculated values and features were stored in struct variables so that VoltView can call them for the ROI explorer plotting and for the on-site analysis.

#### Classification and clustering of o-phys

Parameters for hierarchical clustering were extracted from the o-phys traces by custom-written routines in MATLAB (MathWorks). Parameters were averages of 6-7 sweeps recorded from each ROI. “o-EPSP” and “o-IPSP” was the number of o-PSPs during the 20 Hz Op. “Paired Pulse 2-1” compared the second o-PSP to the first o-PSP amplitude. “Paired Pulse 3-2” compared the third o-PSP to the second o-PSP amplitude. “Subthr Slope q2-q1” compared the mean amplitude of o-Sub between second 250 ms and the first 250 ms. “Subthr Slope q4-q1” compared the mean amplitude of o-Sub between last 250 ms and the first 250 ms. “AP onset” is the delay of the first o-AP during 20 Hz Op. “AP bimodality coeff” calculated Sarle’s bimodality coefficient as the square of skewness divided by the kurtosis, value for uniform distribution is 5/9, values greater than that indicate bimodal (or multimodal) distribution. “AP bimodality binary” gave 1 for bimodal (when coefficient was above 5/9) and 0 for non-bimodal o-AP peak distribution. “AP % in burst” quantified the number of o-APs inside burst periods. “Burst AP freq (Op)” quantified the firing rate inside detected bursts during 20 Hz Op. “AP freq 2nd/1st (Op)” quantified the change of firing rate during the 20 Hz Op comparing the second half (Op-2) with the first half (Op-1). “Burst AP freq (Post)” quantified the firing rate inside detected bursts after 20 Hz Op. “AP freq 2nd/1st (Post)” quantified the change of firing rate after the 20 Hz Op comparing the second half (Post-2) with the first half (Post-1). “AP num” gave the total number of o-APs detected on the baseline of all 6-7 sweeps. “AP freq (Baseline)” was the average firing rate on the baseline before the 20 Hz Op. “AP freq (Op-1)” was the average firing rate on the first half of 20 Hz Op. “AP freq (Op-2)” was the average firing rate on the second half of 20 Hz Op. “AP freq (Post-1)” was the average firing rate on the first half of Post, after 20 Hz Op. “AP freq (Post-2)” was the average firing rate on the second half of Post, after the 20 Hz Op. “Burst num (Op-1)” was the number of burst periods during the first half of 20 Hz Op. “Burst num (Op-2)” was the number of burst periods during the second half of 20 Hz Op. “Burst num (Post-1)” was the number of burst periods during the first half of Post, after the 20 Hz Op. “Burst num (Post-2)” was the number of burst periods during the second half of Post, after the 20 Hz Op. “Burst length (Op-1)” was the average length of burst periods during the first half of 20 Hz Op. “Burst length (Op-2)” was the average length of burst periods during the second half of 20 Hz Op. “Burst length (Op-2/Op-1)” was the change of burst length from Op-1 to Op-2. “Burst length (Post-1)” was the average length of burst periods during the first half of Post, after the 20 Hz Op. “Burst length (Post-2)” was the average length of burst periods during the second half of Post, after the 20 Hz Op. “Burst length (Post-2/Post-1)” was the change of burst length from Post-1 to Post-2.

#### Clustering Analysis

Data analyses were performed in Python 3.8.19 using Visual Studio Code. The following Python packages were utilized during various stages of the analysis: pandas, numpy, seaborn, matplotlib, umap-learn, scikit-learn, scipy, os, io, and re. These tools facilitated tasks ranging from data preprocessing to visualization and clustering.

For clustering, the dataset was scaled using scikit-learn’s StandardScaler to standardize the features. Principal Component Analysis (PCA) was applied to the scaled data, retaining 95% of the total variance to reduce dimensionality while preserving most of the dataset’s information. The preprocessed data was then clustered using scikit-learn’s K-means algorithm, with the number of clusters (k = 13 for AHN, k = 10 for PAG) determined based on the silhouette score and prior knowledge of the dataset. Clustering results were stored as categorical labels in the raw data dataframe, facilitating their integration with downstream analyses and visualizations.

#### Visualization of Clustering Results

To intuitively represent the clustering results, t-distributed Stochastic Neighbor Embedding (t-SNE) was performed. This dimensionality reduction was conducted on the scaled and PCA-transformed data used for clustering. The t-SNE embeddings were computed using the Euclidean distance metric, with dataset-specific perplexity values (47 for AHN and 35 for PAG) optimized for the data structure. The resulting t-SNE coordinates were stored as new columns in the raw data dataframe.

For visualizing specific electrophysiological parameters, numpy’s log1p function was applied directly to the raw data for coloring, without affecting the scaled data or clustering results. This ensured that parameter visualizations accurately reflected their original values, while clustering and t-SNE embeddings were based solely on scaled features.

#### Analysis of Changes After High Frequency Stimulation (HFS)

For both PAG and AHN, cell-level comparisons were conducted between control videos and videos post-high frequency stimulation. Voltview analysis was meticulously configured to prevent frame shifts in cases where cells were not segmented in the HFS videos. Following feature extraction, the comparison was done in Python 3.8.19.

For both datasets, the average firing frequency was computed for each segment following the same methodology as described for the clustering analysis. However, a stricter filtering criterion was applied: frequencies ≤3 were reclassified as 1. To quantify the percentage change induced by HFS, the calculation (HFS − Control) / Control×100, was used. Visualization of these changes was conducted on the t-SNE embedding calculated after the clustering. For color normalization, a centered scale from −100 to 100 was applied to represent changes in a meaningful manner.

#### Post-synaptic response type alignment between AHN-Esr and PAG-Esr datasets

For each dataset (AHN and PAG), the 10 closest samples to the cluster centroids were selected after performing K-means clustering. The mean for each cluster was calculated to define one centroid per cluster. This process was conducted on normalized data using the same feature set specific to each dataset during clustering. Following this, the features were engineered similarly for the collateral somas, normalized using the StandardScaler, and compared with the AHN dataset. Subsequently, pairwise distance metrics from sklearn were used to assign the closest cluster centroid to each cell. This procedure was repeated for the PAG dataset, with the same steps followed for feature engineering, normalization, and centroid assignment.

#### VoltView classifier

We used the unbiased hierarchical clustering of the connectome dataset and calculated the cluster centroids for each o-phys cluster. We had all the 29 parameters in the cluster centroid. We used two parallel classifications and used their consensus. First, for each neuron we calculated the square root of sum squared differences compared to all the cluster centroids and ranked the distances to choose the closest 3 clusters for each neuron. Next for each neuron we calculated the level of correlation to all the cluster centroids, ranked the correlation and choose the 3 highest correlating clusters. If the consensus of the two parallel classifications agreed on one or more clusters, we compared the ranks and assigned the closest cluster to each neuron.

#### Analysis of Allen Transcriptomic Open-Source Data

The Allen Institute published a comprehensive cell-type atlas of the adult mouse brain in 2023, combining single-cell RNA sequencing (sncRNA-seq) and spatial transcriptomics (Yao et al., 2023). Their sncRNA-seq data, generated using two sequencing methods (10Xv2 and 10Xv3), was combined into a unified dataset comprising approximately 4 million cells that passed quality control. For our analysis, we focused on the log2-transformed hypothalamus (HY) dataset generated with the 10Xv2 platform, encompassing approximately 100,000 cells. The data was accessed using the Allen Institute’s publicly available pipeline on GitHub (https://github.com/AllenInstitute/abc_atlas_access/blob/main/descriptions/WMB-10Xv2.md). All data processing and analysis were performed using Python version 3.8.19. To analyze the central and dorsomedial VMH, cells expressing the marker Nr5a1 (also known as SF1) were selected, resulting in a dataset of approximately 2,200 cells. For neuropeptide analysis, the mouse equivalents of the neuropeptide list from the HUGO Gene Nomenclature Committee (HGNC) were utilized, which includes 64 neuropeptides and 4 additional genes encoding tachykinin precursor proteins. Neuropeptides were prioritized based on their mean expression levels, calculated across all SF1 positive cells. Dimensionality reduction was performed on the SF1-expressing cell population by selecting the 100 genes with the highest variance in expression profiles. Additionally, Esr1, Tac1, Pdyn, Gad1, Slc32a1, and Slc17a6 were included due to their relevance to the analysis. Uniform Manifold Approximation and Projection (UMAP) was used for visual dimensionality reduction, with the number of neighbors (n_neighbors) set to 5.

